# Amyloids “at the border”: deep mutagenesis and random sequence extension reveal an incomplete amyloid-forming motif in Bri2 that turns amyloidogenic upon C-terminal extension

**DOI:** 10.1101/2023.09.15.557952

**Authors:** Mariano Martín, Benedetta Bolognesi

**Affiliations:** Institute for Bioengineering of Catalonia (IBEC), The Barcelona Institute of Science and Technology, Baldiri Reixac 10-12, 08028 Barcelona, Spain

## Abstract

Stop-loss mutations cause over twenty different diseases. The effects of stop-loss mutations can have multiple consequences that are, however, hard to predict. Stop-loss in *ITM2B/BRI2* results in C-terminal extension of the encoded protein and, upon furin cleavage, in the production of two 34 amino acid long peptides, ADan and ABri, that accumulate as amyloids in the brains of patients affected by familial Danish and British Dementia. To systematically explore the consequences of Bri2 C-terminal extension, here, we measure amyloid formation for 676 ADan substitutions and identify the region that forms the putative amyloid core of ADan fibrils, located between positions 20 and 26, where stop-loss occurs. Moreover, we measure amyloid formation for ∼18,000 random C-terminal extensions of Bri2 and find that ∼32% of these sequences can nucleate amyloids. We find that the amino acid composition of these nucleating sequences varies with peptide length and that short extensions of 2 specific amino acids (Aliphatics, Aromatics and Cysteines) are sufficient to generate novel amyloid cores. Overall, our results show that the C-terminus of Bri2 contains an incomplete amyloid motif that can turn amyloidogenic upon extension. C-terminal extension with de novo formation of amyloid motifs may thus be a widespread pathogenic mechanism resulting from stop-loss, highlighting the importance of determining the impact of these mutations for other sequences across the genome.

## Introduction

Frameshift mutations and stop-loss mutations make up ∼0.2% of codon-changing mutations^1^ and often result in mRNA degradation by the non-stop mediated-decay (NMD) machinery^2^. However, in the presence of an alternative in-frame stop codon within 50 nt from the original one, NMD is not triggered and aberrant proteins with an extended C-terminus can be expressed^1^. The C-terminal extension resulting from stop-loss can lead to both loss or gain of protein function^3–10^. with over 20 of these mutations proposed as causes of genetic disease^6^.

The molecular mechanisms underlying pathogenicity of stop-loss mutations are diverse^3–10^. The extended protein products resulting from stop-loss can be depleted co- or post-translationally^3,4^. Stop-loss mutations in genes *PNPO, HSD3B2* and *SMAD4* for example produce a novel degron in the extended C-terminal and are targeted to the ubiquitin-proteasome system^5,6^. On the other hand, amorphous aggregates formed by C-terminal extensions of CEP97 are instead cleared by lysosomal degradation^3^. Other mechanisms of stop-loss pathogenicity include mislocalization, with C-terminal extensions causing the transcription factor MITF to mislocalize from the nucleus to the cytoplasm and rhodopsin to occupy different layers of the plasma membrane^7,8^.

Finally, the physico-chemical properties encoded by extended sequences can occasionally lead to aggregation of the mutated protein into amyloid fibrils and thus cause or contribute to the pathogenicity induced by stop-loss mutations. Stop-loss mutations in *NEFH* and *REEP1* allow translation of 3’ UTRs that contain amyloidogenic motifs, leading to protein aggregation and neurodegeneration in Charcott-Marie-Tooth disease and peripheral neuropathy, respectively^9,11^. *ITM2B*, also known as *BRI2*, is instead mutated in familial British (FBD)^10^ and Danish Dementia (FDD)^12^ and also encodes amyloids upon stop-loss. In the last three years, two additional stop-loss mutations in *ITM2B/BRI2* have been discovered, leading to Korean (FKD)^13^ and Chinese Dementia (FCD)^14^. These examples suggest that cryptic amyloid formation could be a more general mechanism of pathogenicity induced by stop-loss and consequent C-terminal extension.

The C-terminus of the transmembrane protein encoded by *ITM2B/BRI2* is cleaved by furin, releasing the 23 amino acid long peptide known as Bri2 (Fig. 1a). Upon stop-loss mutations, furin cleavage produces aberrant 34 amino acid long peptides with distinct amino acid sequences in different types of dementia (Fig. 1b). In FBD, stop-loss results from a single nucleotide variant (SNV), c.799T>A, and produces a peptide known as ABri^10^. FCD and FKD mutations (c.800G>C and c.800G>T, respectively) generate peptides that differ from ABri only in one amino acid at position 24 of the extended peptide (Fig. 1b)^13,14^. In FDD, a 10 nt duplication (c.795-796insTTTAATTTGT) that causes frameshifting and stop loss, leads to an aberrant peptide known as ADan, whose last 12 residues are entirely distinct from ABri (Fig. 1b)^12^. Thus, multiple C-terminal extensions of Bri2 peptide can lead to amyloid formation in the context of autosomal dementia, suggesting that the Bri2 sequence holds an amyloid potential which can be unlocked by specific peptide extensions.

**Figure 1.**
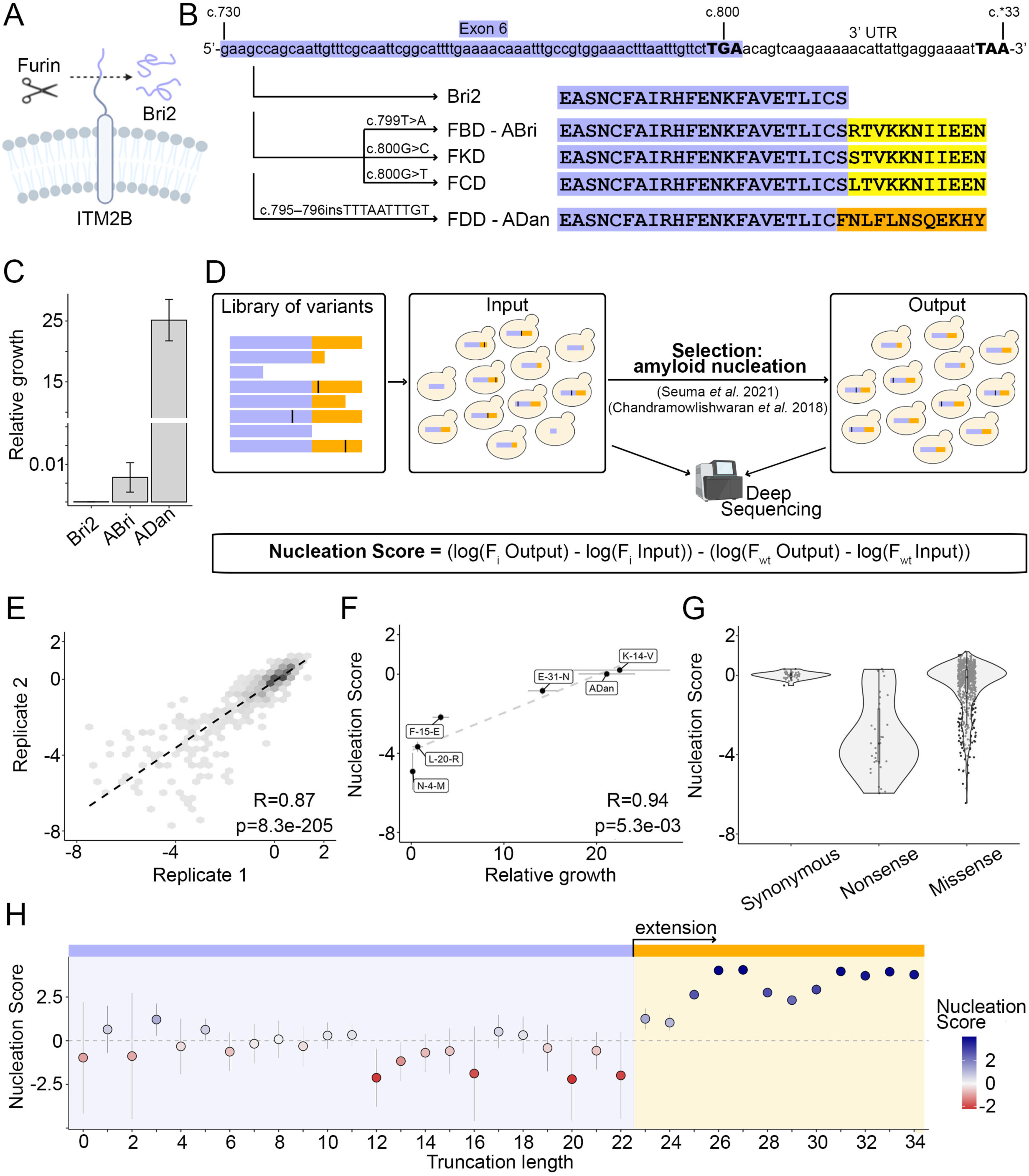
ADan and its truncations are strong nucleators. **A.** Schematics of ITMB2/BRI2 cleavage by the furin protease. **B.** Schematics of *ITMB2/BRI2* and dementia-related mutations at the nucleotide and protein level. cDNA coordinates are detailed at the top and stop codons (conventional or alternative) are represented with bold fonts and upper case. The exon and its translated protein product is highlighted in purple. Extended products resulting from mutations are highlighted in yellow (ABri and ABri-like) and orange (ADan). Familial dementia mutations are detailed over the arrows. **C.** Relative growth of cells expressing Bri2, ABri or ADan fusions calculated as number of colonies growing in the selective conditions of the amyloid nucleation assay (-URA, –Ade) over colonies growing in the absence of selection (-URA)^15^. Error bars indicate the standard deviation of the experiments (n=3). **D.** Deep mutagenesis scheme. A library of sequences of interest is expressed in yeast and selected for amyloid nucleation as reported in Seuma *et al*.^16^. Amyloid nucleation scores are obtained through deep sequencing of input and output samples. **E.** Correlation of nucleation scores between two biological replicates for variants in the ADan library. Pearson correlation coefficient and p-value are indicated. **F.** Correlation between nucleation scores obtained from selection and deep sequencing with relative growth of individual variants in selective over non-selective conditions (n=5). Vertical and horizontal error bars indicate estimated sigma errors and standard deviation of the experiments (n=3), respectively. Pearson correlation coefficient and p-values are indicated. **G.** Distribution of nucleation scores for each class of mutations in ADan library. **H.** ADan truncations nucleation propensity. The horizontal line indicates the weighted mean nucleation score of Bri2 truncations (up to position 22). Vertical error bars indicate 95% confidence interval of the mean.

Here, we employ deep mutational scanning (DMS) to scan the impact of all ADan mutations on amyloid nucleation so to identify the putative amyloid core formed by this peptide. To then explore how likely C-terminal extension in *ITM2B/BRI2* is to generate amyloid cores we use random sequence extension (RSE) and systematically measure the consequences of ∼18,000 random C-terminal extensions of Bri2. Our massive dataset reveals that a large fraction (∼32%) of these extended peptides nucleate amyloids. Among the strongest nucleators we find C-terminal extensions of just two amino acids. We further find that the novel amyloids display specific amino acid enrichments that depend on sequence length.

Altogether, our results suggest that the C-terminus of Bri2 contains an incomplete amyloid motif that can turn amyloidogenic upon extension. They also demonstrate that massively parallel quantitative assays, such as the one presented here, provide a robust strategy to map amyloid formation from novel areas of the sequence space and that they can generate valuable insights into the general consequences of C-terminal extension arising from mutations, drug-induced readthrough or aberrant 3’ UTR translation.

## Results

### Quantifying amyloid nucleation of Bri2 extensions

To measure the ability of Bri2 extensions to nucleate amyloid aggregates we employed a cell-based selection assay where sequences are fused to the nucleation domain of the yeast prion Sup35 (SupN) and amyloid nucleation of the fused peptides seeds the aggregation of endogenous Sup35 (Supp. Fig. 1a)^15^. This process is required for yeast growth in selective conditions and we previously showed that growth rates in this assay correlate with rates of amyloid nucleation in a test tube^16^.

Cells expressing Bri2 do not grow in these conditions, revealing that the unextended peptide does not nucleate amyloid aggregation (Fig. 1c). In sharp contrast, the expression of ADan leads to 25.14 ± 3.41% of the cells growing in selection media over non-selective conditions, a 2-fold increase compared to the growth rate induced in the same conditions by the amyloid-beta peptide (Aβ42), the peptide that forms amyloids in Alzheimer’s disease and is mutated in familial forms of the disease^16^ (Fig. 1c, Supp. Fig. 1b). Cells expressing ABri fusions show very limited growth in the assay (0.0065 ± 0.0039%) with no significant difference compared to expression of the reporter without any fusion (SupN) (Fig. 1c, Supp. Fig. 1b). Growth assays of cells expressing Bri2, ABri or ADan fusions in non-selective conditions revealed that their growth rates were not confounded by differential toxicity (Supp. Fig. 1c).

### Quantifying the effect of ADan substitutions and truncations on amyloid nucleation at scale

To further investigate the sequence-nucleation landscape of the ADan peptide we designed and synthesized a library containing all ADan single amino acid substitutions and truncations (n=680), and expressed it fused to SupN in yeast cells that went under selection as above (Supp. Fig. 1a). We quantified growth rates and therefore amyloid nucleation for each variant in the library by deep sequencing before and after selection (Fig. 1d)^16^. Growth rates, henceforth “nucleation scores”, from this high throughput assay were reproducible between replicates (Fig. 1e, Supp. Fig. 1d) and correlate well with the effects of variants quantified individually (Pearson Correlation, R = 0.94, p-value: 5.3e-3; Fig. 1f). Altogether, we measured amyloid nucleation for 642 single amino acid substitutions (99% of possible amino acid substitutions) and all 34 truncations. In this dataset, synonymous mutations have a nucleation score close to 0, i.e similar to WT ADan, nonsense variants (truncations) have a unimodal distribution with a peak at a nucleation score of ∼-4 and the distribution of missense variants (amino acid substitutions) has a strong bias towards reduced amyloid nucleation (Fig. 1g).

### A four amino acid extension of Bri2 has the same propensity to nucleate amyloids than the full length ADan peptide

Pathogenic Bri2 extensions share the first 22 residues and differ only in the last 12 amino acids (Fig. 1b), suggesting it is the specific extension of ADan that encodes amyloid potential. Measuring the nucleation score of all possible ADan C-terminal truncations in our library, we find that a truncated version of ADan ending at position 25 (equivalent to an extension of Bri2 by F23, N24 and L25) nucleates amyloids significantly faster than Bri2 (Fig. 1h; Z-test, false discovery rate [FDR] = 0.01, p-adjusted: 1.038e-297, Benjamini & Hochberg correction). This effect becomes stronger with a peptide of 26 amino acids (Bri2 extension by F23, N24, L25 and F26), leading to a similar nucleation score as that of the full-length ADan peptide (Z-test, false discovery rate [FDR] = 0.01, p-adjusted: 1.544e-02, Benjamini & Hochberg correction). Together with the previous observation that a 28 residue long ADan truncation forms beta-sheet rich aggregates *in vitro*^17^, these findings suggest that the amyloid potential of ADan is encoded by the first residues that get translated upon stop-loss.

### Deep mutagenesis of ADan uncovers its amyloid core and residues that act as gatekeepers of amyloid nucleation

The distribution of mutational effects for ADan nucleation reveals that 44% of single amino acids substitutions reduce nucleation and only 25% increase it (Z-test, false discovery rate [FDR] = 0.1, Fig. 2a) while 30% of these variants have a nucleation score similar to WT ADan.

**Figure 2.**
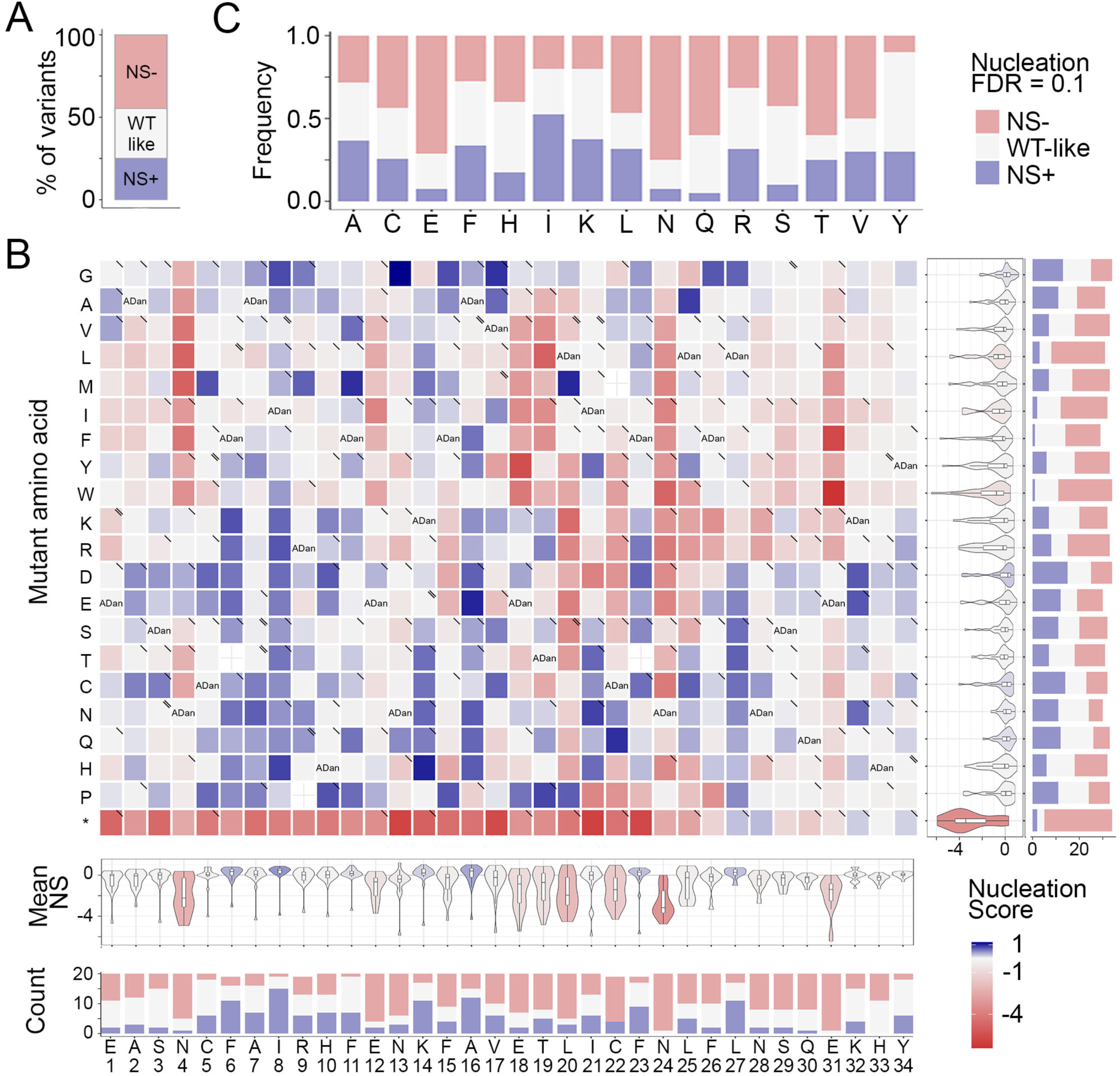
Deep mutagenesis of ADan. **A.** Frequency of variants increasing or decreasing nucleation at FDR=0.1 and variants that are WT-like. **B.** Heatmap of nucleation scores for ADan single amino acid substitutions. The WT amino acid and position are indicated in the x-axis and the mutant amino acid is indicated in the y-axis. Synonymous variants are indicated as “ADan”, missense variants due to SNVs are indicated with “\” and SNVs present in GnomAD with a “\\” in the upper right corner of the cell. The distribution of nucleation scores for each position is summarized in the violin plots below the heatmap. The number of variants increasing or decreasing nucleation at FDR=0.1 per position are indicated as stack bars at the bottom. The distribution of nucleation scores for each mutation is summarized in the violin plots at the right-hand side of the heatmap. The number of variants increasing or decreasing nucleation at FDR=0.1 per mutation are indicated as stack bars at the right. **C.** Frequency of single amino acid substitutions that are increasing or decreasing nucleation (FDR=0.1) upon substituting specific WT amino acid.

Inspecting the heatmap of mutational effects for amino acid changes at all positions in ADan reveals that 26% of the mutations that decrease amyloid nucleation are localized in a short region of 7 amino acids between residues L20-F26 (Fig. 2b, Supp. Fig. 3a). In this continuous stretch of residues most mutations decrease amyloid nucleation (an average of 55% of mutations per position decrease nucleation) and all substitutions to prolines disrupt it (except for L20P that increase nucleation and N24P that is WT-like), suggesting tight structural constraints (Fig. 2b, Supp. Fig. 3a). Thus, we propose that this region forms the inner core of the nucleating ADan fibrils and assign a key role in this process to N24 as all mutations (except N24P) at this position lead to a decrease in nucleation. Charges in this stretch are not tolerated, except at position 23 and change F26E. Outside of this region, at positions E18, T19, N4 and E31, mutations to hydrophobic amino acids decrease nucleation, suggesting longer distance interactions with residues of the amyloid core or a role in modulating the conformations of the monomeric ADan ensemble. Overall, asparagines and glutamic acids are key in ADan nucleation as most substitutions of these amino acids decrease nucleation (75% and 71% of substitutions decrease nucleation, respectively; Fig. 2c).

We identify five gatekeepers, positions where mutations are more likely to increase nucleation than decrease it^18–20^: F6 (12 mutations increasing nucleation and 2 decreasing), I8 (15 increasing), K14 (11 increasing and 2 decreasing), A16 (12 increasing and 5 decreasing) and L27 (11 increasing and 3 decreasing) (Fig.2e). Surprisingly, and in contrast to what was previously found for other amyloids^18–20^, including Aβ42^16,21^, 4 out of 5 gatekeepers are aliphatic residues except for K14.

We also find that 7 of the reported SNPs in *ITM2B/BRI2* (GnomAD database^22^) decrease nucleation, 10 do not affect nucleation and only 1 of them (rs1272422512), which results in mutation R9Q, increases the nucleation of the ADan peptide (Fig. 2b, Supp. Table 1). However, we identify another 53 SNVs that significantly increase nucleation in ADan (Fig. 2b, Supp. Fig. 3a).

### Multiple *ITM2B/BRI2* stop-loss mutations increase nucleation

We then employed the same strategy to test whether specific genetic variants in ABri can boost its weak amyloid nucleation propensity. After data processing and filtering (see methods and Supp. Fig. 2a), we found 121 variants that were able to increase nucleation (Supp. Fig. 2 and Supp. Fig. 3b). The heatmap of mutational effects for amino acid changes at all positions in ABri reveals that substitutions R24L and R24S increase nucleation (Supp. Fig. 2d and Supp. Fig. 3b), meaning that the peptides that are a product of FCD and FKD mutations are more prone to aggregate than both Bri2 and ABri. Moreover, the R24C mutation, equivalent to a single stop-loss mutation leading to a Cys codon (c.801A>C: TGA>TGC or c.801A>T: TGA>TGT), significantly increases nucleation (Supp. Fig. 2b,d and Supp. Fig. 3b). Also, substitution by Cys at position 20 (L20C) significantly increases nucleation, inducing a relative growth in selection media over non-selective conditions of 63.1 ± 7.1%, that is 2.7 times more than ADan peptide nucleation (23.1 ± 3.6%).

### Different types of mutations increase the nucleation of ADan, ABri and Aβ42 amyloids

As Alzheimer’s disease and *ITM2B/BRI2*-related familial dementias share common clinical and neuropathological features, including amyloid depositions^12,23–25^, we evaluated if the impact of mutations on amyloid nucleation of ADan depends on similar properties as those driving Aβ42 nucleation. There is no significant correlation between the nucleation scores of variants in these peptides when comparing mutations of the same amino acids (Fig. 3a) or mutations to the same amino acid (Fig. 3b). In addition, to compare mutational effects by position, we aligned ADan and Aβ42 peptides in three different ways, at the N-terminal, C-terminal and with ClustalO^26^. Regardless of the strategy, no significant correlation at the aligned positions was detected (Supp. Fig. 4a,b,c).

**Figure 3.**
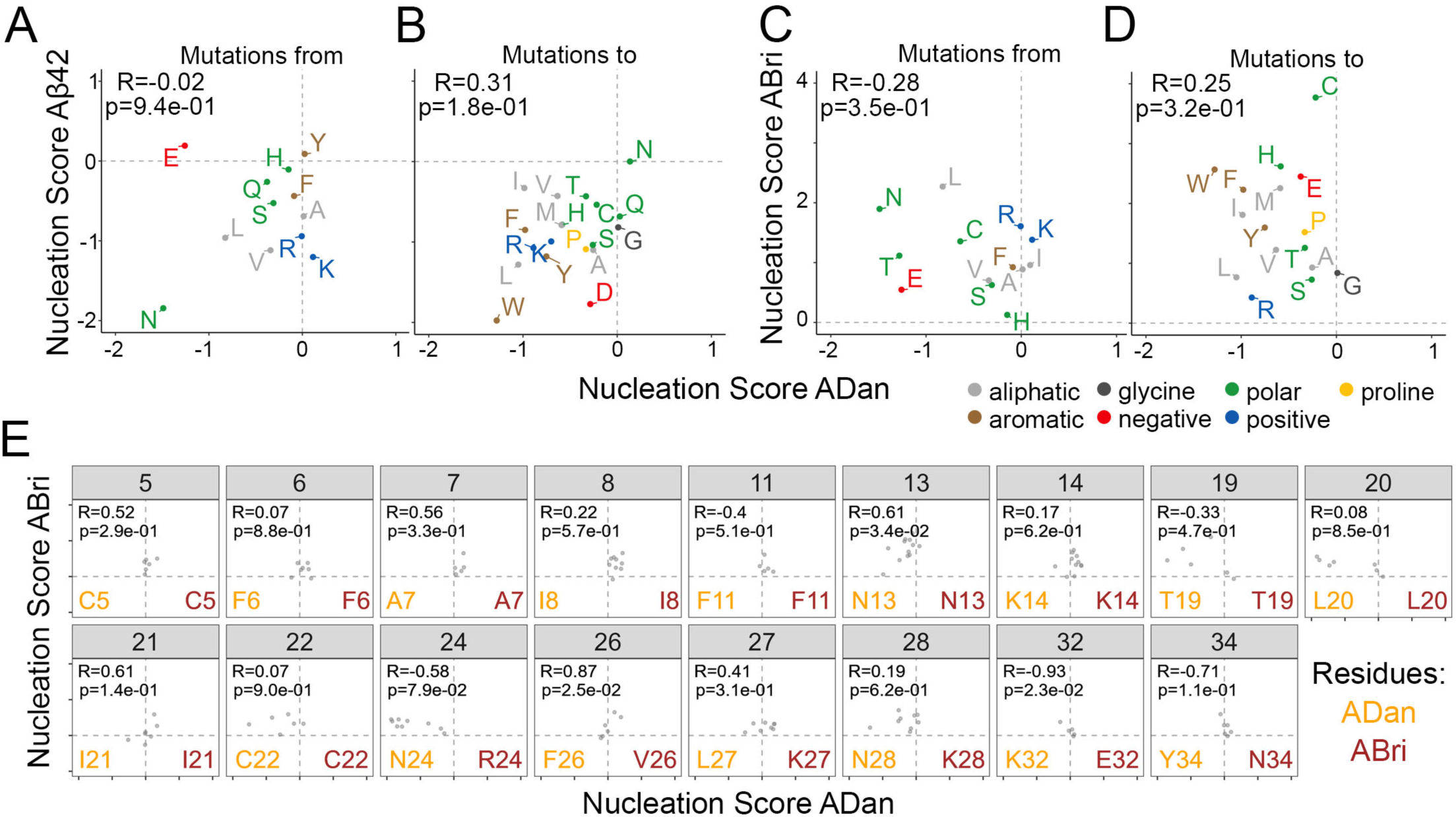
Comparing the mutational effects of single AA variants in ADan, Aβ42 and ABri libraries. **A-B.** Correlation of average nucleation scores for each type of substitution of the same amino acid (**A**) or to the same amino acid (**B**) in ADan and Aβ42. **C-D.** Correlation of average nucleation scores for each substitution to the same amino acid (**C**) or of the same amino acid (**D**) in ADan and ABri. **E.** Correlation of ADan and ABri nucleation scores for positions where more than 4 ABri variants were measured. ADan residues are detailed in orange and ABri residues in red. Pearson correlation coefficient and p-value are indicated in each plot.

Similarly, we compared ADan and ABri nucleation scores. None of the comparisons between ADan and ABri revealed significant correlations between the nucleation scores of these peptides (Fig. 3c,d; Supp. Fig. 4d). We also tested the correlation in those positions with at least 5 confident measurements in the ABri library, uncovering significant positive correlations only for positions 13 (N), 26 (F and V) and 32 (K and E, ADan and ABri respectively; Fig. 3E).

Surprisingly, changes in hydrophobicity can explain very little of the variance in the ADan dataset (Supp. Fig. 5a). Moreover, state-of-the-art amyloid predictors scores poorly correlate with ADan nucleation scores (Supp. Fig. 5a) and analysis of the net and total charge of ADan variants gives little insight in the process of ADan amyloid nucleation (Supp. Fig. 5b).

### Bri2 random extensions reveal the properties of de novo amyloid nucleating peptides

To better understand the sequence determinants of amyloid core generation upon Bri2 C-terminal extension, we generated three libraries where the Bri2 peptide (22 amino acids) is extended by 12 random degenerate (NNK, where N = A/C/G/T and K = G/T) codons, in a way that is similar to what happens in ADan biogenesis (ADan-like extensions, Fig. 4A). Each library was selected independently, and sequencing was used to quantify the relative amyloid nucleation scores for a total of 17,952 unique amino acid sequences corresponding to 4.68e-18 of the possible sequence space (20^12^) (Fig. 4b and Supp. Fig. 6a). The β-sheet propensity and hydrophobicity, relevant features in amyloid nucleation, of this experimental sequence space overlap with those of the human proteome (Fig. 4d). After data processing and quality control, the vast majority of sequences had a nucleation score of 0. Consequently, we classified sequences with a nucleation score significantly greater than 0 as nucleators (n=5,678; 31.6% of the sequences; Z-test, false discovery rate [FDR] = 0.05), and all other sequences as non-nucleators (n=12,274; 68.4%; Fig. 4b). The resulting nucleation scores correlate well with the effect of sequences quantified individually (Fig. 4c).

**Figure 4.**
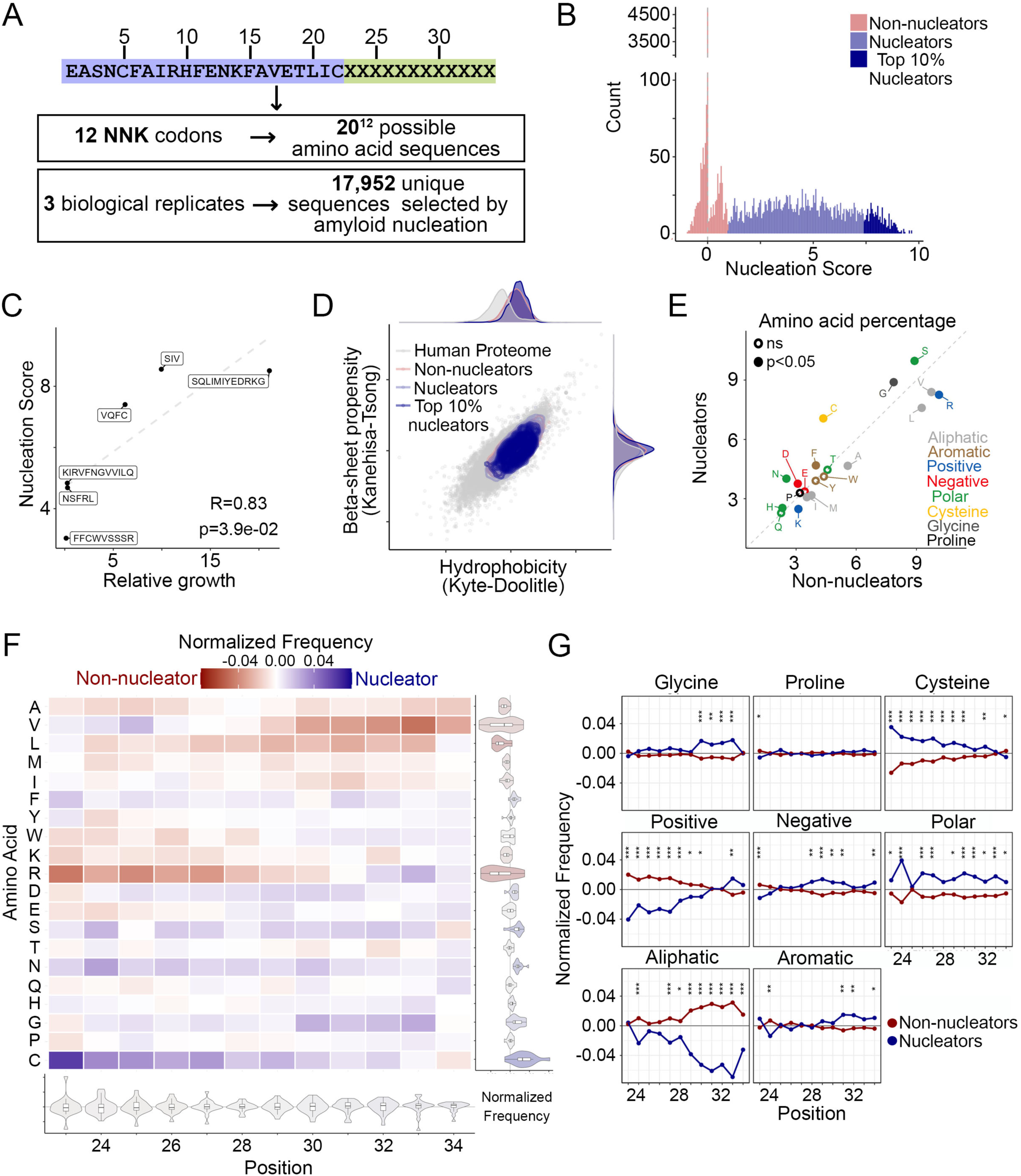
Amyloid nucleation by Bri2 random extensions of 12 amino acids. **A.** Random extensions design and sequence space. A library of random sequences encoded by 12 degenerated NNK codons was expressed in yeast and selected for amyloid nucleation. **B.** Nucleation scores distribution of random sequences present in replicate 1. Variants are classified as non-nucleators (red), nucleators (blue) or top 10% nucleators (dark blue). **C.** Correlation between nucleation scores obtained from selection and deep sequencing with relative growth of individual variants in selective over non-selective conditions (n=6). Pearson correlation coefficient and p-values are indicated. **D.** Hydrophobicity (Kyte-Doolitle^41^) and beta-sheet propensity (Kanehisa-Tsong^42^) of assayed sequences relative to human proteome. **E.** The percent composition of residues grouped by their physicochemical properties in nucleators and non-nucleators sequences. **F.** Heatmap of normalized frequencies for Bri2 random extensions. The distribution of normalized frequencies for each position is summarized in the violin plots below the heatmap and the distribution of normalized frequencies for each mutation is summarized in the violin plots at the right-hand side of the heatmap. **G.** The position-specific differences in amino acid type frequencies across nucleating and non-nucleating sequences. Asterisks indicate marginal p-value (chi-square test). p<0.05: *; p<0.01: **; p<0.001: ***.

We examined the differences in amino acid frequency between nucleating and non-nucleating sequences with an extension of 12 amino acids (Fig. 4e). Although the differences were modest, several of these were statistically significant (t-test), owing to the large sample size of our data. The most relevant differences are in Asn (p-value=4.28e-56, Cohen’s d effect size=0.30, difference in frequency=1.50), Cys (p-value=6.69e-110, effect size=0.43, difference=2.71) and Arg (p-value=5.23e-30, effect size=0.22, difference=-1.87; Fig. 4e; Supp. Fig. 6c; Supp. Table 2). We also analyzed the amino-acid composition position-wise, finding that towards the N-terminus of the extension (close to the end of the Bri2 sequence) nucleators are significantly enriched (chi-squared test) in polar residues (Fig. 4f,g; min. p-value position 24=3.53e-11, Cohen’s d effect size=0.25, difference in frequency=0.056), Cys, Phe and/or Val (Fig. 4f, Supp. Fig. 6c; Supp. Table 3). They are instead depleted in charged residues and aliphatics (Fig. 4f,g; Supp. Table 4). The enrichment in polars and Cys is maintained all along the whole extension while the depletion in aliphatics becomes even stronger towards the C-terminus (Fig. 4f,g; Supp. Table 4). In addition, at the C-terminus, nucleators are enriched in Gly (Fig. 4f,g; min. p-value position 33=1.54e-5, effect size=0), aromatics as well as positive and negative charges (Fig. 4f,g; Supp. Table 4).

Among the random C-terminal extensions, 1,824 sequences (∼10% of the total) have a Ser residue at position 23, mimicking the effect of stop-loss mutations that lead to production of ABri and effectively generating a subset of ABri-like extensions. We performed the same amino acid enrichment analysis on this subset of extensions, finding similar results as the ones found for all the other random extensions (Supp. Fig. 7).

We next evaluated if common amyloid aggregation predictors are able to predict the nucleation of these sequences and if they can discriminate between non-nucleatores and nucleators (Supp. Fig. 8a), or between non-nucleators and the 10% strongest nucleators in each random library (Supp. Fig. 8b). All of the predictors show a poor or limited performance on the prediction task (Supp. Fig. 8). In both cases CamSol outperforms the rest of the predictors with an AUC of 0.5660±0.0086 for discriminating non-nucleators versus nucleators and an AUC of 0.6482±0.0154 for non-nucleators versus the 10% strongest nucleators.

### Bri2 extensions of two specific amino acids are sufficient to nucleate amyloids

Our random libraries also included shorter C-terminal extensions due to premature stop codons in the 12 degenerate-codon sequence (Fig. 5a; number of sequences with an extension < 12 amino acids: 5,110). We analyzed these sequences and found that the resulting peptide length is not correlated with the strength of amyloid nucleation (Fig. 5b). Moreover, we found that extending Bri2 by only two specific amino acids can create strong nucleating peptides among the 10% strongest nucleators in the dataset (Fig. 5c,e). We analyzed the position-specific differences between nucleators and non-nucleators composition and found that the amino acid composition of C-terminal extensions with length 8-11 amino acids (peptides length of 30-33 residues) is similar to that of the 12 amino acid C-terminal extensions (Supp. Fig. 9). However, for C-terminal extensions of length <8 amino acids (peptides length of 24-29 residues), nucleators exhibit a significant enrichment in aromatics at the N-terminus (Fig. 5d and Supp. Fig. 9), revealing that amyloid nucleation is promoted by distinct types of residues depending on peptide length. In ABri-like extensions (i.e. sequences with a Ser at position 23), we find that a C-terminal extension of two specific amino acids, where at least one of them is aliphatic, creates strong nucleators (Fig. 5E).

**Figure 5.**
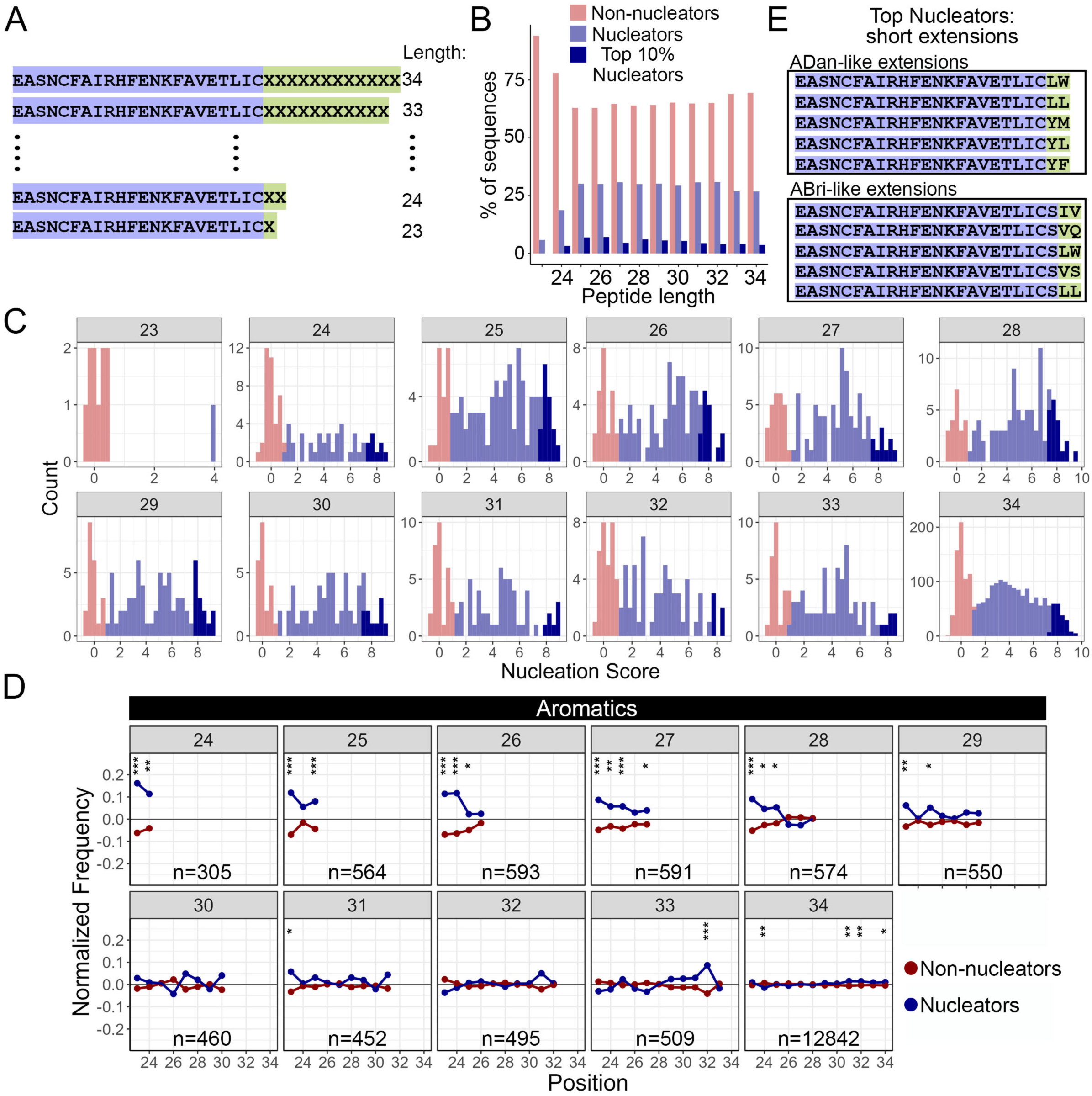
Nucleators resulting from Bri2 random extensions of different lengths have different specific amino acid enrichment. **A.** Range of lengths for the Bri2 random extensions measured in the assay. **B.** Percentage of non-nucleators, nucleators and top 10% nucleators sequences for each peptide length. **C.** Nucleation scores distribution of random extended sequences present in replicate 1. The length of the peptide is indicated inside the grey box of each plot. Variants are classified as non-nucleators (red), nucleators (blue) or top 10% nucleators (dark blue). **D.** The position-specific differences in aromatic amino acids frequencies across nucleating and non-nucleating sequences in truncations of different length. The length of the peptide is indicated inside the grey box of each plot. Asterisks indicate marginal p-value (chi-square test). p<0.05: *; p<0.01: **; p<0.001: ***. **E.** Top nucleating sequences of ADan-like extensions (top) and ABri-like extensions (bottom) of 2 amino acids. Bri2 sequence is highlighted in purple and the extended sequence in green.

## Discussion

*ITM2B/BRI2* mutations affecting its stop codon are associated with familial forms of dementia and deposits containing amyloid forms of the peptides which are generated upon stop-loss have been observed *in vivo*^10,12,27^. Here, we systematically addressed how sequences resulting from stop-loss in *ITM2B/BRI2* can nucleate amyloids using deep and random mutagenesis combined to a selection assay that reports on the amyloid nucleation of thousands of sequences of different length in parallel.

The ADan peptide very rapidly forms amyloids in this cell-based assay, 2-fold faster than Aβ42 for reference, but ABri is at the lower end of the detection capability of our assay. While this sharp difference could be due to limitations in this specific assay, it is in line with results in transgenic mice and drosophila models expressing ABri, which do not exhibit amyloid deposits^28^ and feature low neurotoxicity, in contrast with those expressing ADan^27,29^.

Thanks to this massively parallel assay, we identify a continuous stretch of residues (L20-F26) in ADan which are crucial for amyloid nucleation and likely constitute the core of ADan amyloids. Surprisingly, changes in hydrophobicity can explain very little of the variance in the ADan dataset (Supp. Fig. 5). It would have been impossible to predict the mutational impact in ADan on the basis of simple physicochemical properties, aggregation predictors or even the latest generation of variant effect predictors^30^. What is more, it would have also been impossible to predict mutational impact in ADan on the basis of the complete mutational landscape of another extracellular amyloid peptide of similar length, Aβ42^21^, suggesting that our understanding of the process of amyloid formation by novel peptides is still data limited and highlighting the need of systematically measuring amyloid nucleation for these and other peptides that form amyloids and lead to neurodegeneration^31,32^.

By employing unbiased random sequence extension, we expanded the probed sequence space both in terms of length and identity, quantifying amyloid nucleation for ∼18,000 sequences. To our knowledge, this is the largest data set of protein sequence extensions currently available.

∼32% of them are able to nucleate amyloids. This dataset also reveals that a C-terminal amino acid extension of just two residues is enough to significantly increase the propensity of Bri2 to nucleate amyloids. Other short C-terminal extensions (peptide length < 34 amino acids) also generate strong nucleators. These results are in agreement with a previous study showing that an ADan truncation at position 28 can form beta-sheet rich aggregates *in vitro*^17^. They also suggest that an ‘incomplete amyloid motif’ exists at the end of the Bri2 sequence, and that specific C-terminal extensions upon stop loss or due to insertions before the stop codon can turn it into an effective driver of amyloid nucleation.

Cryptic amyloid motifs have been discovered in the 3’ UTRs of at least three genes that are mutated in neurodegenerative disease^9–11^. Our datasets can now contribute to build better models that can identify and predict more of these motifs. In addition, while stop-codon readthrough of TGA codons happens at a low frequency of 0.01-0.1%^33,34^ this phenomenon can be promoted by small molecules^35^ such as Gentamicin or ELX-02, which are under clinical trial for the treatment of Herlitz junctional epidermolysis bullosa and Cystic Fibrosis, respectively (clinical trials: NCT04140786 and NCT04135495). Along this line, the possibility of quantifying the impact of C-terminal sequence extensions provides a systematic way to uncover incomplete or hidden amyloids across the genome, gaining the ability to preventively interpreting the impact of stop-codon readthrough on de-novo amyloid formation.

## Acknowledgements

Work in the lab of BB is supported by the la Caixa Research Foundation project ‘DeepAmyloids’ (LCF/PR/HR21/52410004), by the Spanish Ministry of Science, Innovation and Universities (PID2021-127761OB-I00 and RYC2020-028861-I, funded by MCIN/AEI/10.13039/501100011033, “ERDF A way of making Europe” and “ESF Investing in your future”). and by the European Union (ERC Consolidator, Glam-MAP, 101125484). Views and opinions expressed are however those of the author(s) only and do not necessarily reflect those of the European Union or the European Research Council. Neither the European Union nor the granting authority can be held responsible for them. IBEC is a member of the CERCA Program / Generalitat de Catalunya. We thank Ben Lehner and Mike Thompson for discussing the random extension dataset. We thank the Chernoff lab for providing strains and plasmids and the CRG Genomics core technology for sequencing. Figure 1a was created with BioRender.com.

## Data availability

Raw sequencing reads and processed nucleation scores are deposited in NCBI’s Gene Expression Omnibus (GEO) as GSE244612 and GSE270792. The processed reads and nucleation scores are also available as Supplementary Data 1.

## Code availability

All scripts used for downstream analysis and to reproduce all figures are in https://github.com/BEBlab/ADan-BriNNK.

## Methods

### Library design

The designed libraries contain a total of 1088 unique ADan nucleotide variants (680 single protein variants encoded by 1, 2, and 3 nucleotide changes) and 1088 unique ABri nucleotides variants (680 single protein variants encoded by 1, 2, and 3 nucleotide changes). The random libraries were designed with the sequence encoding for the Bri2 peptide (22 amino acids) at the N-terminus followed by random extensions of 12 amino acids encoded by degenerate codons (NNK, where N = A/C/G/T and K = G/T).

### Plasmid Library construction

Libraries were synthesized by Integrated DNA Technologies (IDT) as oligo pools covering the ADan and/or ABri 102 nt sequence or as an ultramer of 12 NNK codons for the random libraries (36 nucleotides), flanked by 25 nt upstream and 21 nt downstream constant regions for cloning. 2 ul of 10uM library pool were extended into a double stranded DNA by a single cycle PCR (Q5 high-fidelity DNA polymerase, NEB) with primers annealing to the constant regions (primers MM_01, Supplementary Data 2). The product was purified from a 2% agarose gel (QIAquick Gel Extraction Kit, Qiagen). In parallel, the pCUP1-Sup35N plasmid was linearized by PCR (Q5 high-fidelity DNA polymerase, NEB; primers MM_02-04, Supplementary Data 2). The product was purified from a 1% agarose gel (QIAquick Gel Extraction Kit, Qiagen). The oligo pool was then ligated into 200 ng of the linearized plasmid in a 1:10 (vector:insert) ratio by a Gibson approach with 3 h of incubation at 50°C followed by dialysis for 3 h on a membrane filter (MF-Millipore 0.025 μm membrane, Merck) and vacuum concentration. The product was transformed into 10-beta Electrocompetent *E. coli* (NEB), by electroporation at 2.0 kV, 200 Ω, 25 μF (BioRad GenePulser machine). For the random libraries, the electroporation was performed three times independently to obtain three random libraries. Cells were recovered in SOC medium for 30 min and grown overnight in 30 ml of LB ampicillin medium. A small amount of cells were also plated in LB ampicillin plates to assess transformation efficiency. A total of >1M transformants were estimated, meaning that each variant in the ADan and ABri libraries are represented ∼1000 times. For the three random libraries, ∼5M, ∼8M and ∼2M transformants were obtained, respectively. 5 ml of overnight cultures were harvested to purify the ADan and ABri libraries with a mini prep (QIAprep Miniprep Kit, Qiagen). 50 ml of overnight culture were harvested to purify each random library with a midi prep (Plasmid MIDI Kit, Qiagen).

### Yeast transformation

*Saccharomyces cerevisiae* [psi-pin-] (MATα ade1–14 his3 leu2-3,112 lys2 trp1 ura3–52) provided by the Chernoff lab^15^ was used in all experiments in this study. Yeast cells were transformed with the ADan and ABri libraries in three biological replicates. Per replicate, an individual colony was grown overnight in 3 ml YPDA medium at 30 °C and 4 g. Cells were diluted in 40 ml to OD600 = 0.3 and grown for 4–5 h. When cells reached the exponential phase (OD∼0.8–0.9) cells were harvested at 3000 × g for 5 min, washed with 50 ml milliQ, centrifuged at 3000 × g for 5 min and washed with 25 ml SORB buffer (100 mM LiOAc, 10 mM Tris pH 8.0, 1 mM EDTA, 1 M sorbitol). Cells were resuspended in 1.4 ml of SORB and incubated 30 min on an orbital shaker. After incubation, 800 ng of library and 30 ul of ssDNA (UltraPure, Thermo Scientific) were added and incubated for 5 min at room temperature and then 10 min on an orbital shaker. 6 ml of YTB-PEG (100 mM LiOAc, 10 mM Tris pH 8.0, 1 mM EDTA, 40% PEG 3350) and 580 ul of DMSO were added to the cells. Heat-shock was performed at 42 °C for 20 min in a liquid bath with intermittent shaking. Cells were harvested and incubated in 50 ml of recovery medium (YPDA medium + Sorbitol 0.5 M) for 1 h at 30 °C. Cells were harvested and grown in 50 ml plasmid selection medium (-URA, 2% glucose) for 50 h at 30 °C. A small amount of cells were also plated in plasmid selection solid medium to assess transformation efficiency. 60,000-100,000 transformants were estimated for each biological replicate, meaning that each variant in the library is represented at least 50 times. After 50 h, cells were diluted in 50 ml plasmid selection medium to OD= 0.05 and grown exponentially for 15 h. Finally, the culture was harvested and stored at −80 °C in 25 % glycerol.

### Large-scale yeast transformation of random libraries

Yeast cells were transformed with the plasmid random libraries midi prep in three independent experiments. For each of the experiments, an overnight pre-growth culture in 25 ml of YPDA medium at 30°C was diluted to OD600 = 0.3 in 175 ml YPDA and incubated at 30°C 200 rpm for ∼4 hr. When cells reached the exponential phase, they were harvested, washed with milliQ, and resuspended in sorbitol mixture (100 mM LiOAc, 10 mM Tris pH 8, 1 mM EDTA, 1M sorbitol). After a 30 min incubation at room temperature (RT), 4 ug of plasmid library and 175 µl of ssDNA (UltraPure, Thermo Scientific) were added to the cells. YTB-PEG mixture (100 mM LiOAc, 10 mM Tris pH 8, 1 mM EDTA pH 8, 40% PEG3350) was also added and cells were incubated for 30 min at RT and heat-shocked for 15 min at 42°C in a water bath. Cells were harvested, washed, resuspended in 250 ml recovery medium (YPD, sorbitol 0.5M, 70 mg/L adenine) and incubated for 1 hr at 30°C 200 rpm. After recovery, cells were resuspended in 350 ml -URA plasmid selection medium and allowed to grow for 50 hr. Transformation efficiency was calculated for each tube of transformation by plating an aliquot of cells in -URA plates. Each of the three random libraries was bottlenecked to ∼500,000 yeast transformants. Two days after transformation, the culture was diluted to OD600 = 0.08 in 500 ml -URA medium and grown until exponential phase. At this stage, cells were harvested and stored at -80°C in 25% glycerol.

### Selection experiments

*In vivo* selection assays were performed in three independent biological replicates for ADan and ABri libraries. For each replicate, cells were thawed from −80 °C in 50 ml plasmid selection medium at OD = 0.05 and grown until exponential for 15 h. At this stage, cells were harvested and resuspended in 50 ml protein induction medium (-URA, 2% glucose, 100 μM Cu_2_SO_4_) at OD = 0.1. After 24 h the 40 ml input pellets were collected, and cells were plated on -ADE-URA selection medium in 145-cm2 plates (Nunc, Thermo Scientific). Plates were incubated at 30 °C for 7 days. Finally, colonies were scraped off the plates with PBS 1x and harvested by centrifugation to collect the output pellets. Both input and output pellets were stored at −20 °C before DNA extraction. Three input and three output samples were processed for sequencing. For each of the three random libraries, one selection experiment was performed, and one input sample and three technical replicates of the output pellet were processed for sequencing as described above.

### DNA extraction and sequencing library preparation

Input and output pellets were resuspended in 0.5ml extraction buffer (2% Triton-X, 1% SDS, 100 mM NaCl, 10 mM Tris-HCl pH 8, 1 mM EDTA pH 8). They were then frozen for 10 min in an ethanol-dry ice bath and heated for 10 min at 62 °C. This cycle was repeated twice. 0.5ml of phenol:chloroform:isoamyl (25:24:1 mixture, Thermo Scientific) was added together with glass beads (Sigma). Samples were vortexed for 10 min and centrifuged for 30 min at 2000 × g. The aqueous phase was then transferred to a new tube, and mixed again with phenol:chloroform:isoamyl, vortexed and centrifuged for 45 min at 2000 × g. Next, the aqueous phase was transferred to another tube with 1:10 V 3 M NaOAc and 2.2 V cold ethanol 96% for DNA precipitation. After 30 min at −20 °C, samples were centrifuged and pellets were dried overnight. The following day, pellets were resuspended in 0.3 ml TE 1X buffer and treated with 10 μl RNAse A (Thermo Scientific) for 30 min at 37 °C. DNA was finally purified using 10 μl of silica beads (QIAEX II Gel Extraction Kit, Qiagen) and eluted in 30 μl elution buffer. Plasmid concentrations were measured by quantitative PCR with SYBR green (Merck) and primers annealing to the origin of replication site of the PCUP1-Sup35N plasmid at 58 °C for 40 cycles (primers MM_05-06, Supplementary Data 2). The library for high-throughput sequencing was prepared in a two-step PCR (Q5 high-fidelity DNA polymerase, NEB). In PCR1, 50 million of molecules (ADan and ABri libraries) or 300 million molecules (random libraries) were amplified for 15 cycles with frame-shifted primers with homology to Illumina sequencing primers (primers MM_07–26, Supplementary Data 2). The products were treated with ExoSAP treatment (Affymetrix) and purified by column purification (MinElute PCR Purification Kit, Qiagen). They were then amplified for 12 cycles in PCR2 with Illumina-indexed primers (primers MM_27–67, Supplementary Data 2). For ADan and ABri libraries, the six samples (one input-output pair per biological replicate) were pooled together equimolarly and the final product was purified from a 2% agarose gel with 20 μl silica beads (QIAEX II Gel Extraction Kit, Qiagen). The three random libraries were purified individually. The libraries were sent for 125 bp paired-end sequencing in an Illumina HiSeq2500 sequencer at the CRG Genomics core facility. In total, >10 million paired-end reads were obtained for each of ADan and ABri libraries, representing >1000x read coverage. Each of the random libraries obtained >45 million paired-end reads.

### Individual variant testing

Selected variants, from ADan and ABri libraries, for individual testing were obtained by PCR linearisation (Q5 high-fidelity DNA polymerase, NEB) with mutagenic primers (primers MM_68-85, Supplementary Data 2) or by ultramer amplification and Gibson assembly with the linearized plasmid for the sequences coming from the random libraries (primers MM_86-91). PCR products were treated with Dpn1 overnight and transformed in DH5ɑ competent *E. coli*. Plasmids were purified by miniprep (QIAprep Miniprep Kit, Qiagen) and transformed into yeast cells. All mutated plasmids were verified by Sanger sequencing. For measuring growth in selective conditions, yeast cells expressing individual variants were grown overnight in a plasmid selection medium (-URA 2% glucose). They were then diluted to OD 0.1 in protein induction medium (-URA 2% glucose 100 μM Cu_2_SO_4_) and grown for 24 h. Cells were plated on -URA (control) and -ADE-URA (selection) plates in three independent replicates and allowed to grow for 7 days at 30 °C. Adenine growth was calculated as the percentage of colonies in -ADE-URA relative to colonies in -URA.

For individual growth rate measurements, yeast cells expressing individual variants were grown overnight in plasmid selection medium (-URA 2% glucose) and diluted to OD 0.2 until exponential. They were then diluted again to OD 0.1 in non-inducing (-URA 2% glucose) and inducing (-URA 2% glucose 100 μM Cu_2_SO_4_) protein expression mediums. Cell growth was measured at 30 °C for >48 h at 10 min intervals in a microplate reader (Spark, Tecan) in three biological replicates. Growth rates were calculated using the GrowthCurver package in R.

### Data processing

FastQ files from paired end sequencing of the libraries were processed using DiMSum (https://github.com/lehner-lab/DiMSum)^36^, an R pipeline for analyzing deep mutational scanning data. 5′ and 3′ constant regions were trimmed, allowing a maximum of 20% of mismatches relative to the reference sequence. Sequences with a Phred base quality score below 30 were discarded. Non-designed variants were also discarded for further analysis, as well as variants with fewer than 100 (random libraries) or 200 (ADan and ABri libraries) input reads in all of the replicates.

### Nucleation scores and error estimates

The DiMSum package (https://github.com/lehner-lab/DiMSum)^36^ was also used to calculate nucleation scores (NS) and their error estimates for each variant in each biological replicate as:

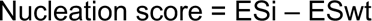

Where ESi = log(Fi OUTPUT) - log(Fi INPUT) for a specific variant and ESwt = log(Fwt OUTPUT) - log(Fwt INPUT) for ADan or ABri WT.

NSs for each variant were merged across biological replicates using error-weighted mean and centered to the WT NS. All NS and associated error estimates are available in Supplementary Data 1.

The ADan library has a good signal/noise ratio due to its significant growth in the selection assay. After NSs centering, each variant was tested against the WT NS at FDR=0.1 and classified in three possible groups: WT-like, NS_inc and NS_dec. Sigma values were normalized to the interquartile range and WT-like variants with a normalized sigma value above a cut-off of 0.2 were excluded. The same procedure was part of ABri library processing. As this library has a poor signal/noise ratio (Supp. Fig. 2a), few variants passed the filtering process. NS was obtained for 676 unique ADan variants and 679 unique ABri variants. 664 confident estimates (with low normalized sigma value) were obtained for the ADan library and 177 in the ABri library.

For the random extensions libraries, 1 input and 3 output sequencing results were used. DimSum was run using an arbitrary WT sequence chosen among those with high input counts. The sequences used were: TTGTAGGTTGCTCAGGAGGTTAATACGTATACGTAG, TTTTGTTAGTTTGATGTGCATGAGTGTCTGTAGTGT and TGGAAGGGGACGATGGTTGTGGGTAGTAATTGGCCG for experiment 1, 2 and 3, respectively.

NSs for each sequence were merged within each replicate using error-weighted mean and centered to the mode value of the distribution (as most of the sequences do not nucleate). Those sequences that are present in the input and not in the output were imputed with the mode value of the distribution. Most of the sequences have a nucleation score of 0. Consequently, we classified sequences with a nucleation score significantly (FDR=0.05) greater than 0 as nucleators and all other sequences as non-nucleators. Duplicated sequences were merged and mean nucleation score and the mode of the nucleator status was calculated. We discarded those non-nucleators sequences that had big errors and a mean nucleation score above the minimum nucleator score. For each of the random libraries we classified the top 10% nucleator sequences as another group. Finally, variants from the three experiments were merged and duplicated variants were treated as previously described. 17,952 unique sequences with NS were obtained (7,783 experiment 1; 5,562 experiment 2 and 4,607 experiment 3). All NS estimates are available in Supplementary Data 1.

### Aggregation predictors

For the aggregation predictors (Tango, Amypred, Camsol, Aggrescan)^37–40^ scores were calculated per sequence and, if necessary, individual residue level scores were summed to obtain a score per single AA mutation sequence. We also used the Kyte-Doolittle hydrophobicity scale^41^.

### Statistics and reproducibility

Based on the transformation efficiency, each variant in the designed libraries (n = 1088 nucleotide variants each) is expected to be represented at least 10x at each step-in selection experiments and library preparation. In terms of sequencing, reads that did not pass the QC filters using the DiMSum package were excluded (https://github.com/lehner-lab/DiMSum)^36^. The experiments were not randomized. The Investigators were not blinded to allocation during experiments and outcome assessment.

---

**Supplementary Figure 1.**
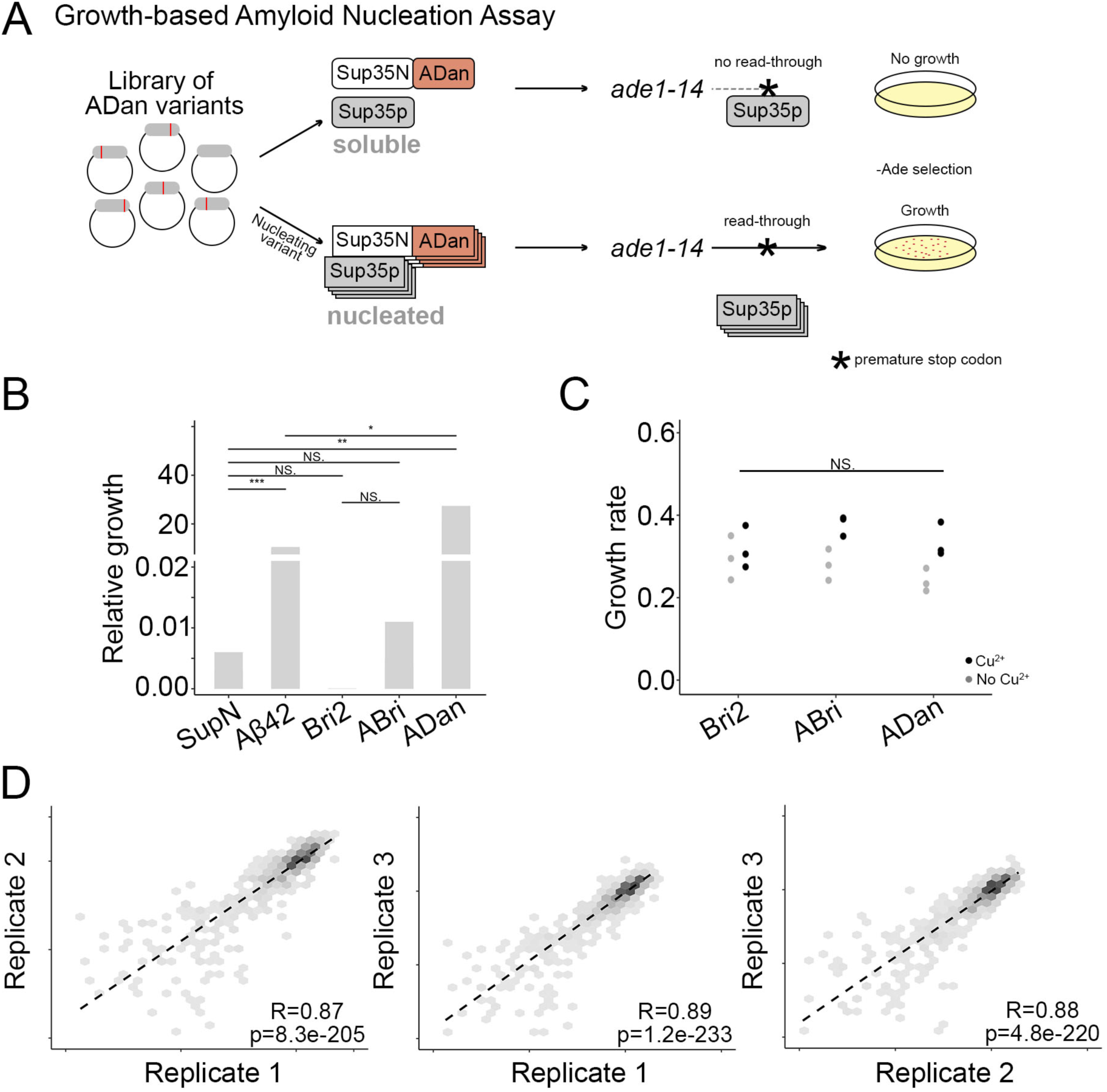
ADan and ABri nucleation. **A.** Schematics of the reporter assay. **B.** Individual testing of SupN (negative control), Aβ42 (positive control), Bri2, ABri and ADan in the reporter assay. Relative growth of cells expressing each sequence, calculated as the number of colonies growing in the selective conditions of the amyloid nucleation assay (-URA, –Ade) over colonies growing in the absence of selection (-URA). t-test significance is indicated (n=3 biological replicates/variant). **C.** Individually measured growth rates for Bri2, ABri and ADan, in non-inducing (no Cu^2+^) and inducing (Cu^2+^) protein expression conditions (n=3 biological replicates/variant). One-way ANOVA with Dunnett’s multiple comparisons test against Bri2 Cu^2+^. **D.** Correlation of nucleation scores between biological replicates for variants in the ADan library. Pearson correlation coefficient and p-value are indicated.

**Supplementary Figure 2.**
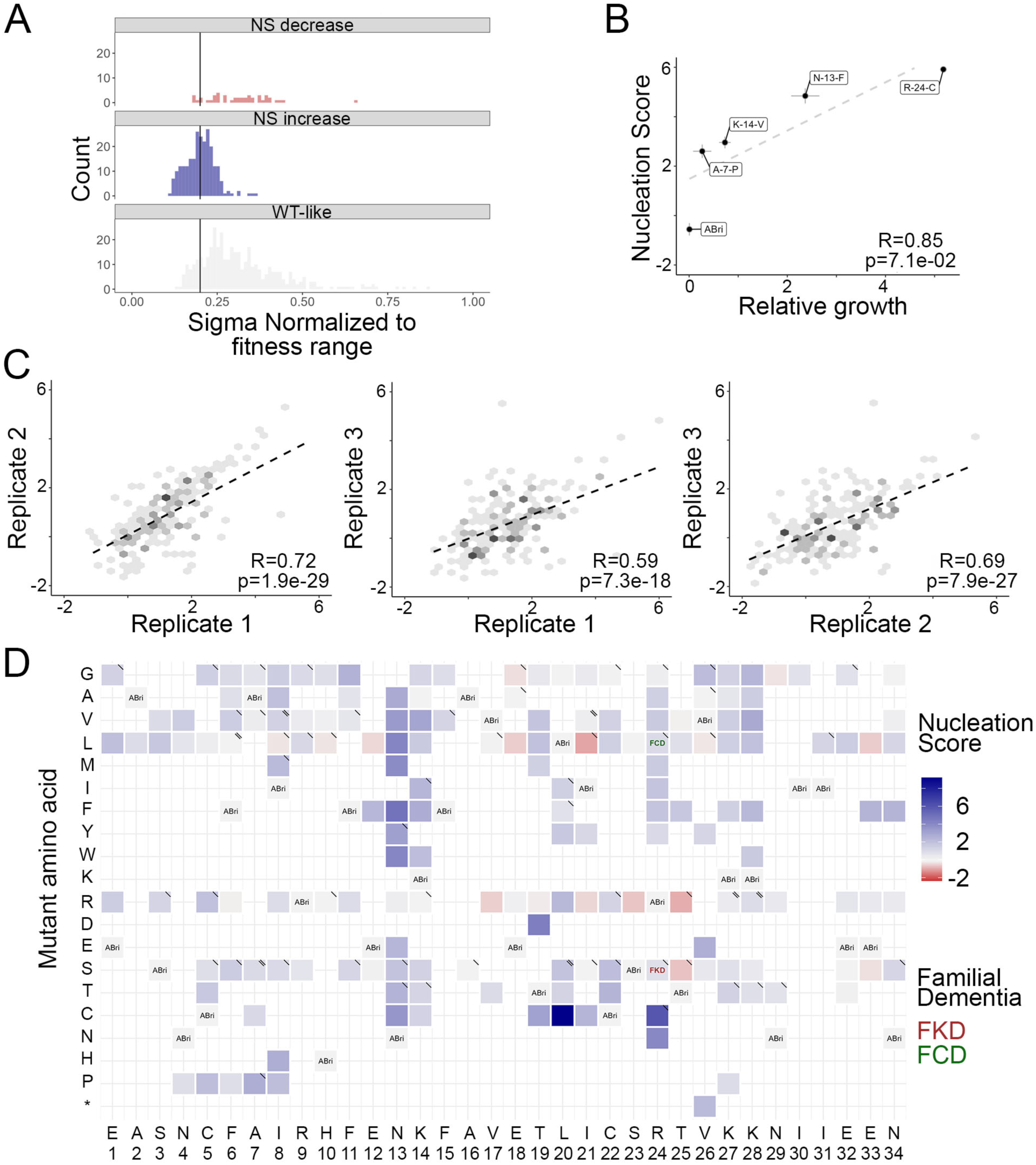
Deep mutagenesis of ABri. **A.** Normalized sigma distribution for variants increasing, decreasing or not affecting nucleation at FDR=0.1. Vertical line indicates threshold value for considering low sigma. **A.** Correlation of nucleation scores between two biological replicates for variants in the ABri library. Pearson correlation coefficient and p-value are indicated. **B.** Correlation of nucleation scores obtained from selection and deep sequencing with relative growth of individual variants in selective conditions (n=5). Vertical and horizontal error bars indicate estimated sigma errors and standard deviation of the experiments (n=3), respectively. Pearson correlation coefficient and p-values are indicated. **C.** Correlation of nucleation scores between biological replicates for variants in the ABri library. Pearson correlation coefficient and p-value are indicated. **D.** Heatmap of nucleation scores for ABri single amino acid substitutions. The WT amino acid and position are indicated in the x-axis and the mutant amino acid is indicated in the y-axis. Synonymous variants are indicated as “ABri”, missense variants due to SNVs are indicated with “\” and SNVs present in GnomAD with a “\\” in the upper right corner of the cell. Missing variants have been excluded on the basis of poor sequencing quality and low normalized sigma values (see Methods).

**Supplementary Figure 3.**
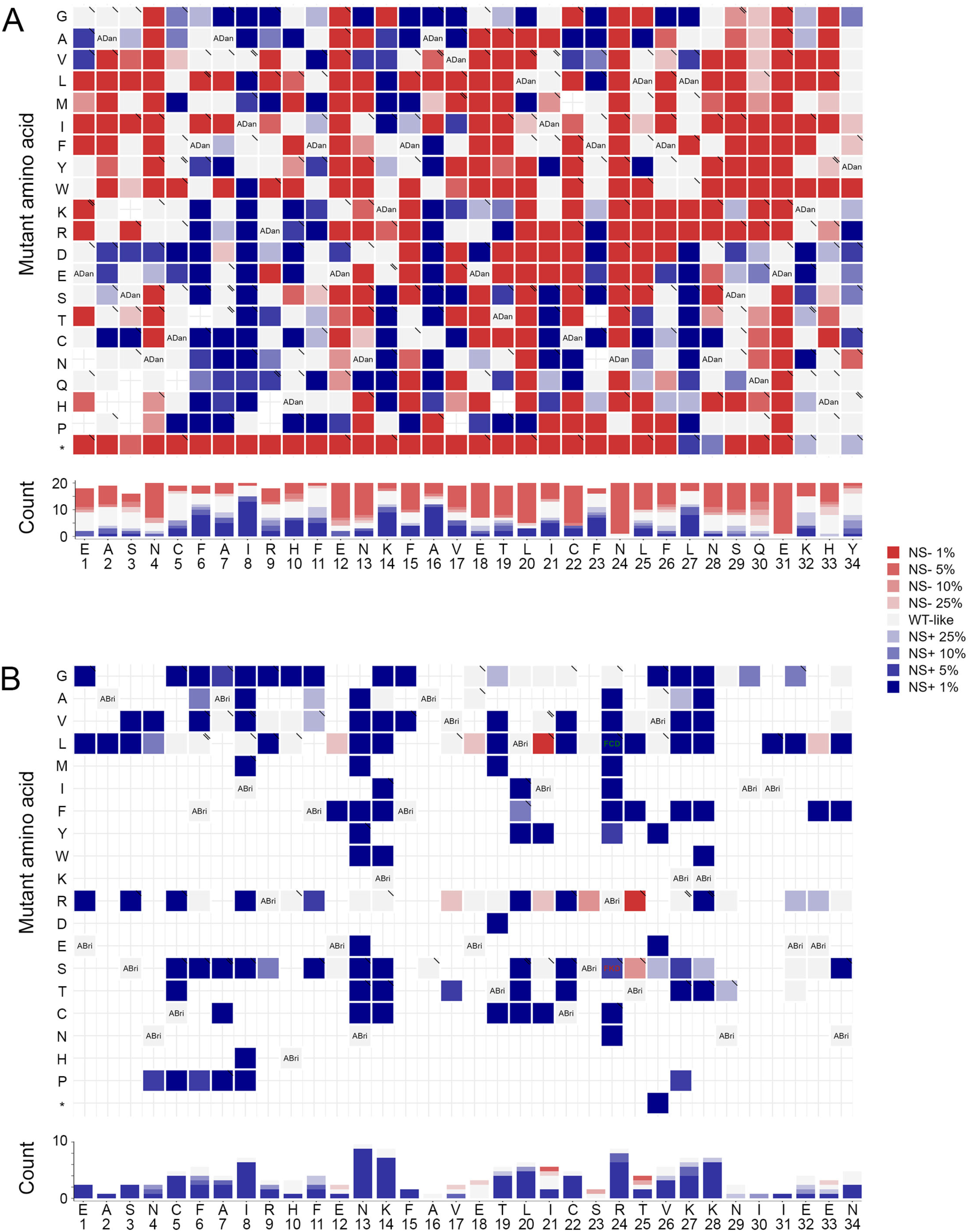
Mutational effects of single AA substitutions. Heatmap of FDR categories for the effects of single amino acid substitutions. The WT amino acid and position are indicated in the x-axis and the mutant amino acid is indicated in the y-axis. Synonymous mutants are indicated with “ADan” (**A**) or “ABri” (**B**). Missense variants due to SNVs present in GnomAD are indicated with “\” in the upper right corner of the cell. The number of variants increasing or decreasing nucleation at different FDRs per position are indicated as stack bars at the bottom.

**Supplementary Figure 4.**
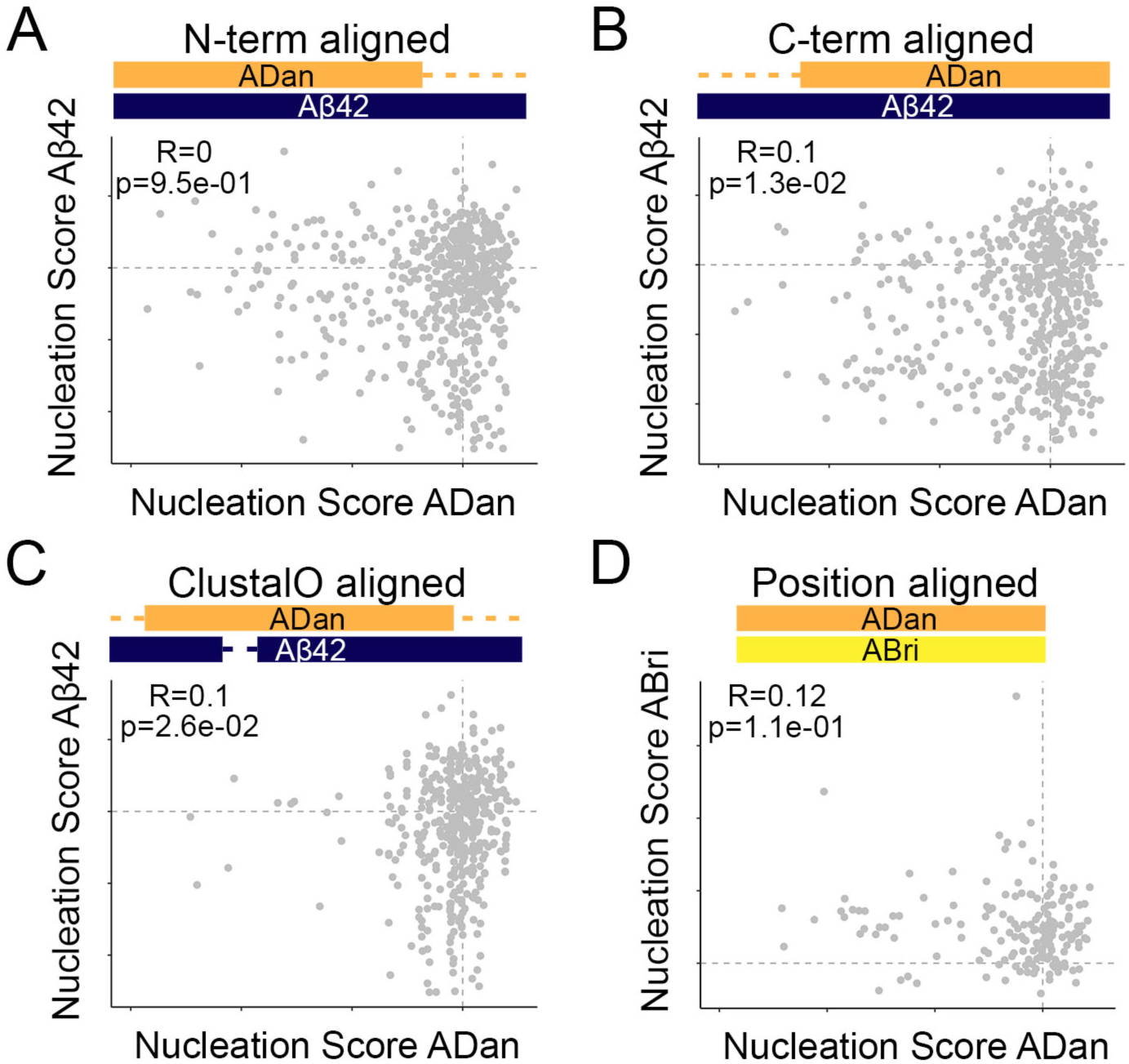
Comparing the mutational effects of single amino acid variants in ADan, Aβ42 and ABri libraries. Correlation of nucleation scores between ADan and Aβ42 aligned at N-term (**A**), at C-term (**B**) and with ClustalO (**C**), and between ADan and ABri (**D**).

**Supplementary Figure 5.**
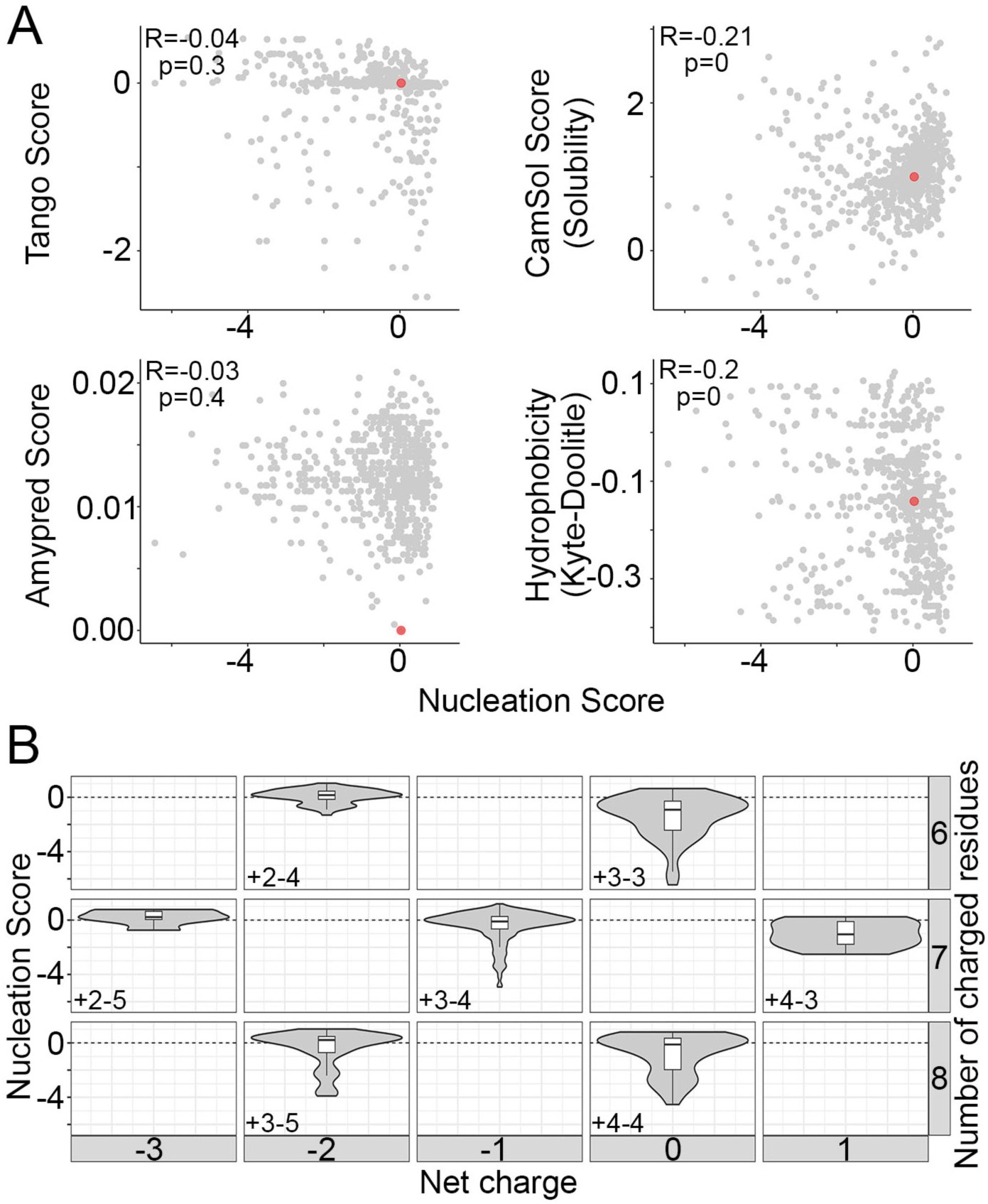
Physicochemical features partially explain nucleation. **A.** ADan nucleation scores correlation with amyloid propensity predictors (Tango and Amypred), solubility predictors (CamSol) and Kyte-Doolittle hydrophobicity score. Red dots indicate the values for the WT sequence. Pearson correlation coefficient and p-values are indicated. **B.** Nucleation score distributions arranged by the number of charged residues (y-axis) and the total net charge (x-axis) in ADan. Numbers inside each cell indicate the number of positive and negative residues. The horizontal line indicates WT nucleation score (0).

**Supplementary Figure 6.**
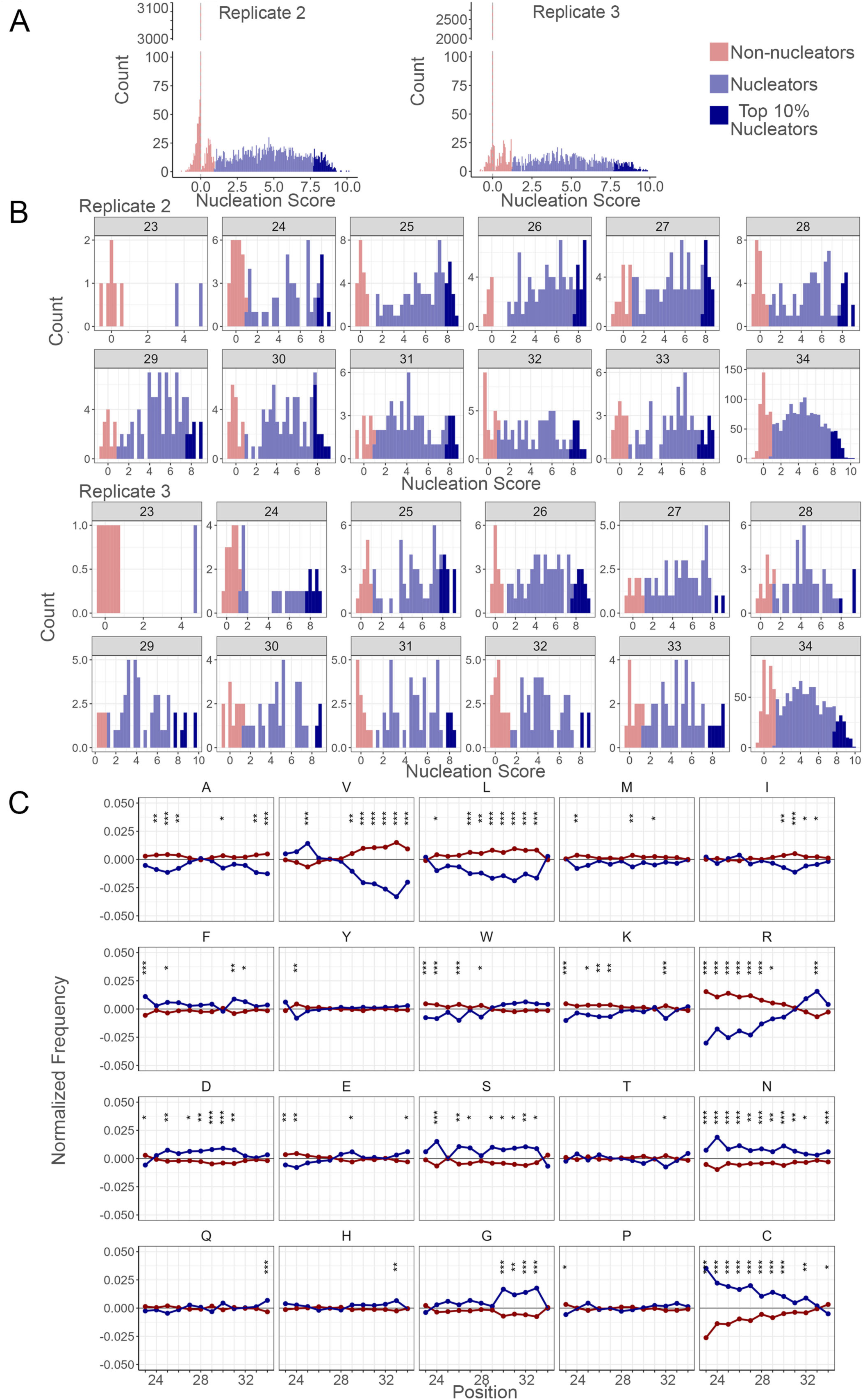
Bri2 random extensions. **A.** Distribution of nucleation scores for random sequences of length 34 for replicates 2 and 3. Variants are classified as non-nucleators (red), nucleators (blue) or top 10% nucleators (dark blue). **B.** Nucleation scores distribution of random sequences present in replicate 2 and 3 (top and bottom, respectively). The length of the peptide is indicated inside the grey box of each plot. Variants are classified as non-nucleators (red), nucleators (blue) or top 10% nucleators (dark blue). **C.** The position-specific differences in amino acid frequencies of sequences 34 amino acid long, across nucleating and non-nucleating sequences. Asterisks indicate marginal p-value (chi-square test). p<0.05: *; p<0.01: **; p<0.001: ***.

**Supplementary Figure 7.**
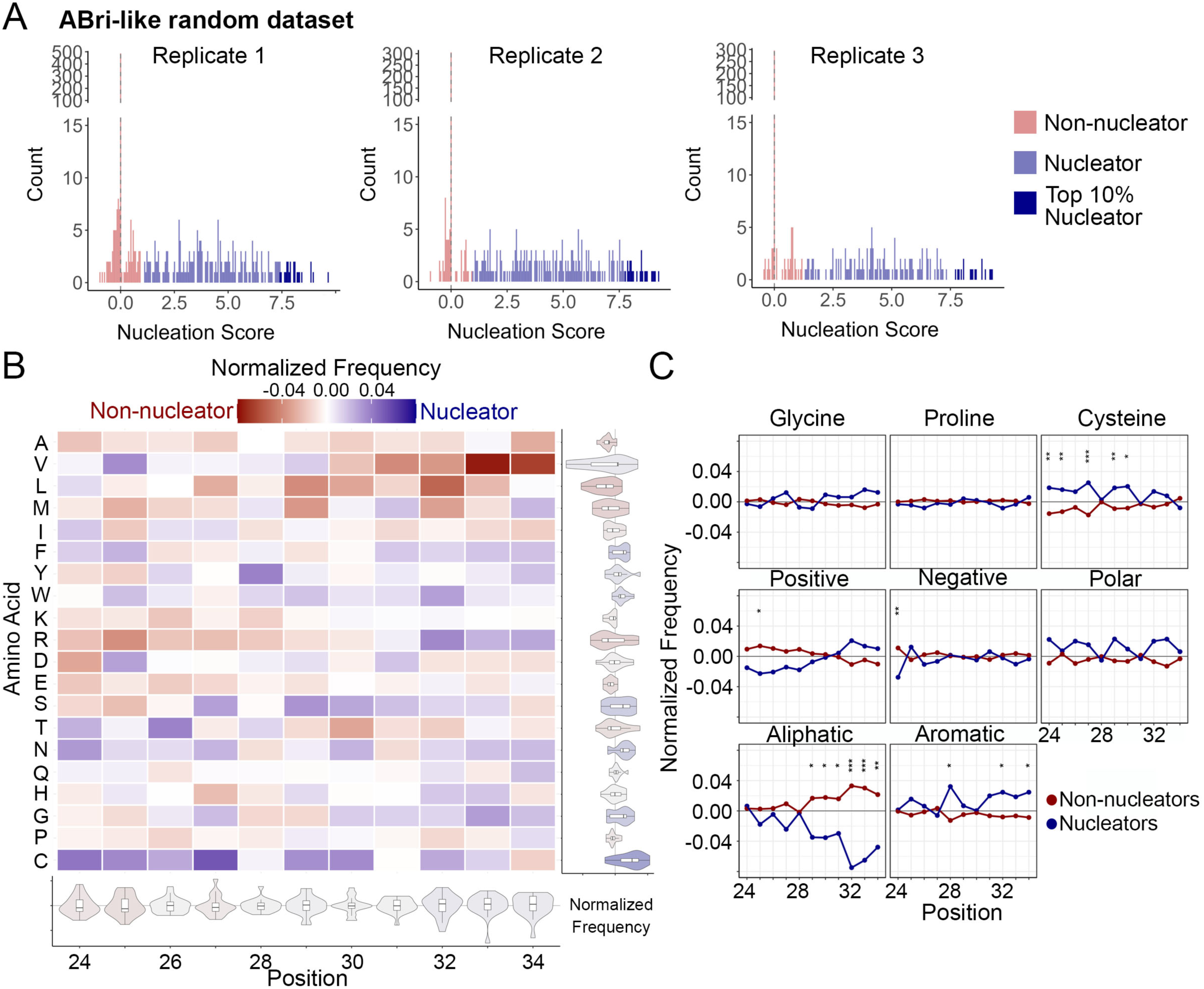
ABri-like random dataset. **A.** Nucleation scores distribution of random sequences present in the three replicates. Variants are classified as non-nucleators (red), nucleators (blue) or top 10 nucleators (dark blue). **B.** Heatmap of normalized frequencies for ABri-like random extensions. The distribution of normalized frequencies for each position is summarized in the violin plots below the heatmap and the distribution of normalized frequencies for each mutation is summarized in the violin plots at the right-hand side of the heatmap. **C.** The position-specific differences in amino acid type frequencies across nucleating and non-nucleating sequences. Asterisks indicate marginal p-value (chi-square test). p<0.05: *; p<0.01: **; p<0.001: ***.

**Supplementary Figure 8.**
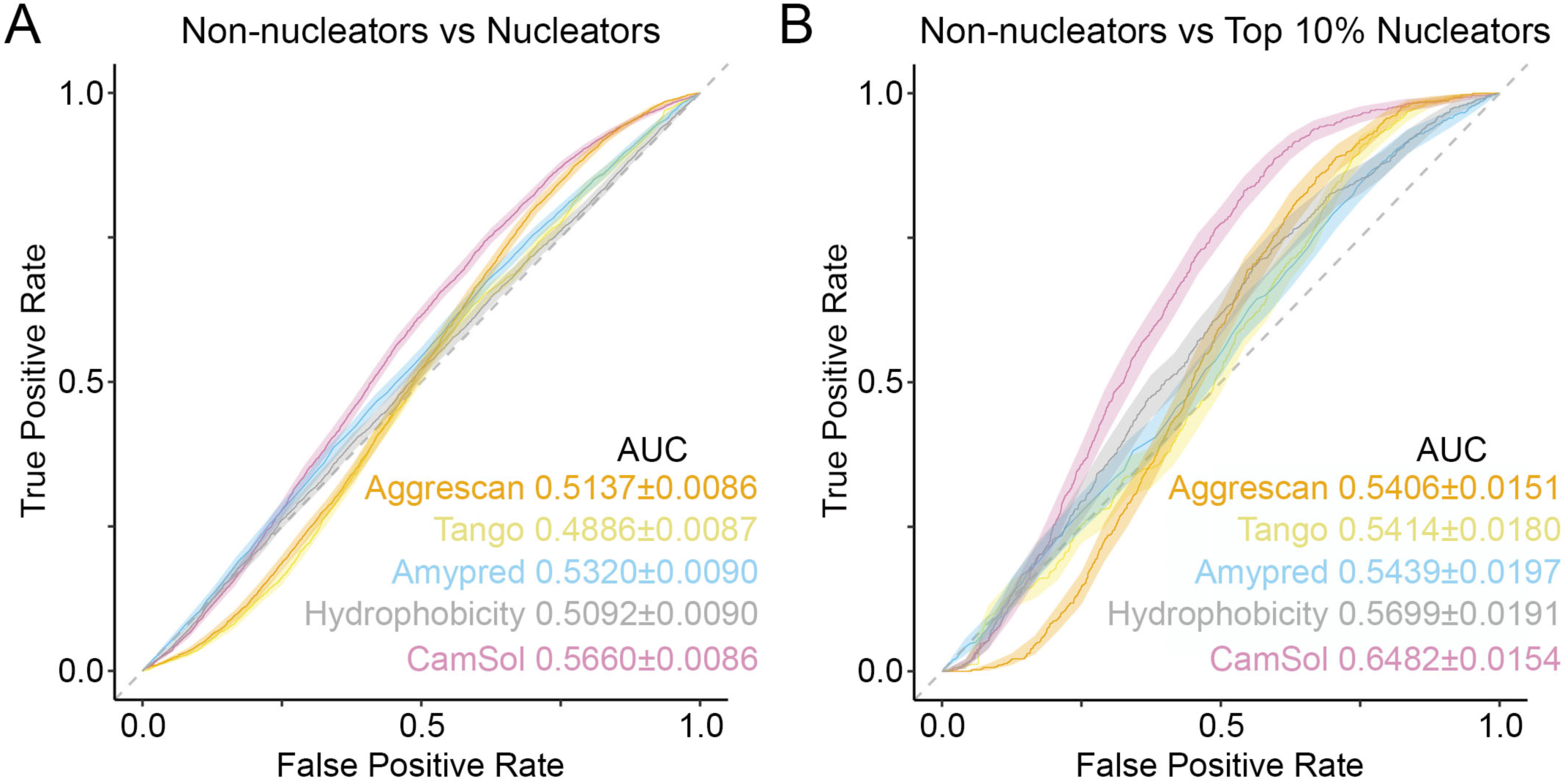
Performance of amyloid predictors on Bri2 random extension dataset. ROC curves of state-of-the-art amyloid predictors on non-nucleators vs nucleators (**A**) and non-nucleators vs top 10% nucleators sequences (**B**). AUC and confidence interval is detailed for each of the predictors.

**Supplementary Figure 9.**
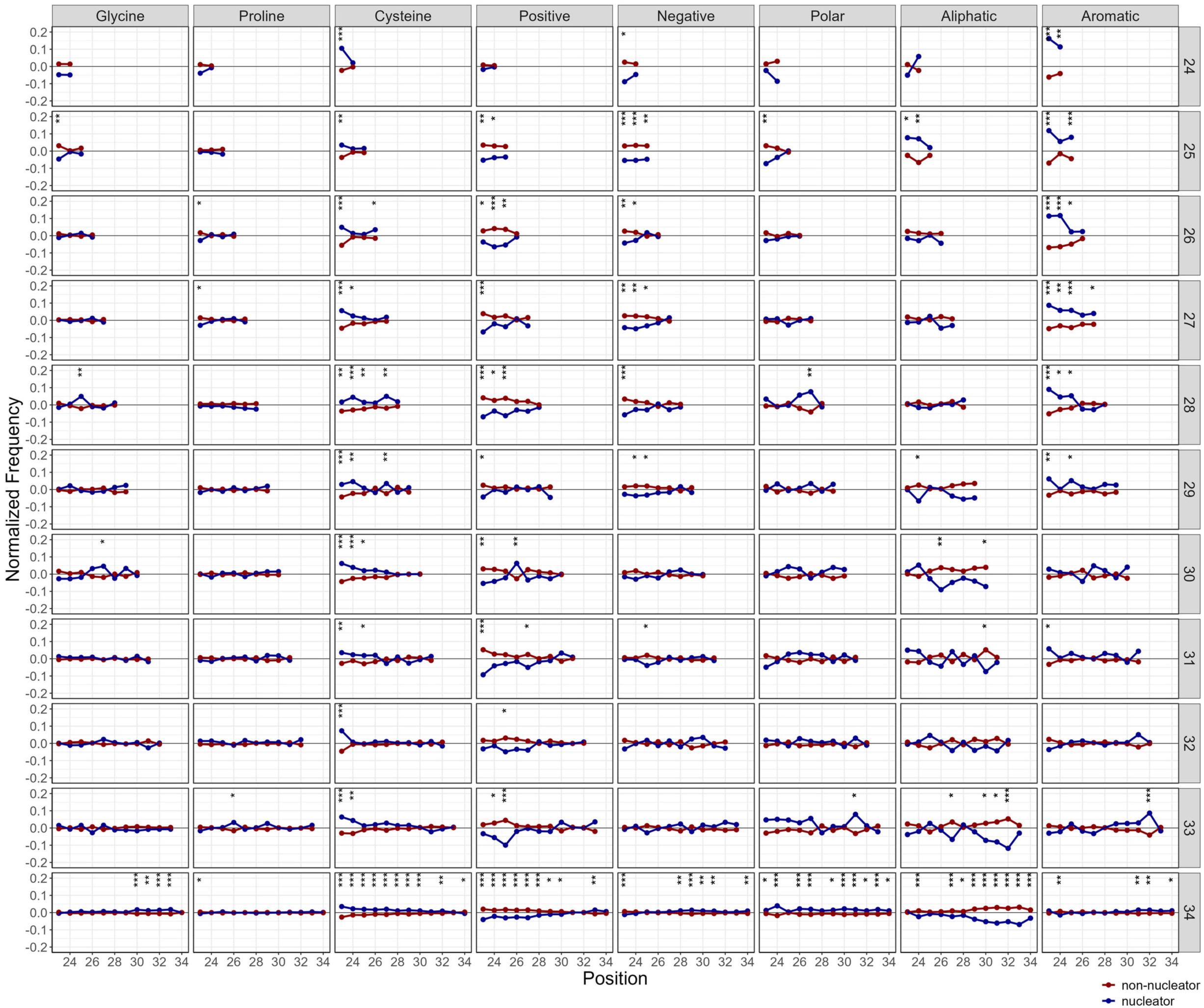
Position-specific differences in amino acid type frequencies for truncated sequences across nucleating and non-nucleating sequences. Asterisks indicate marginal p-value (chi-square test). p<0.05: *; p<0.01: **; p<0.001: ***.

